# Sample preparation for tissue mass spectrometry-based (phospho)proteomics

**DOI:** 10.64898/2026.06.17.732915

**Authors:** Sibilla Sander, Ibrahim Bayramoglu, Michael Stumpe, Gaetana Restivo, Mitchell P. Levesque, Joern Dengjel

## Abstract

This protocol describes the workflow for the preparation of tissue samples for proteome and phosphoproteome analyses using mass spectrometry. The tissue samples are cryogenically pulverized and homogenized in a sucrose-based buffer to ensure proper tissue disruption. For depletion of lipid contaminants, proteins are purified using chloroform-methanol precipitation, followed by a resuspension in a urea-based buffer for enzymatic digestion. Peptides are desalted and enriched for phosphopeptides prior LC-MS/MS analysis. The workflow was developed for skin biopsies but is compatible with a broad range of tissue types.

**Graphical abstract:** 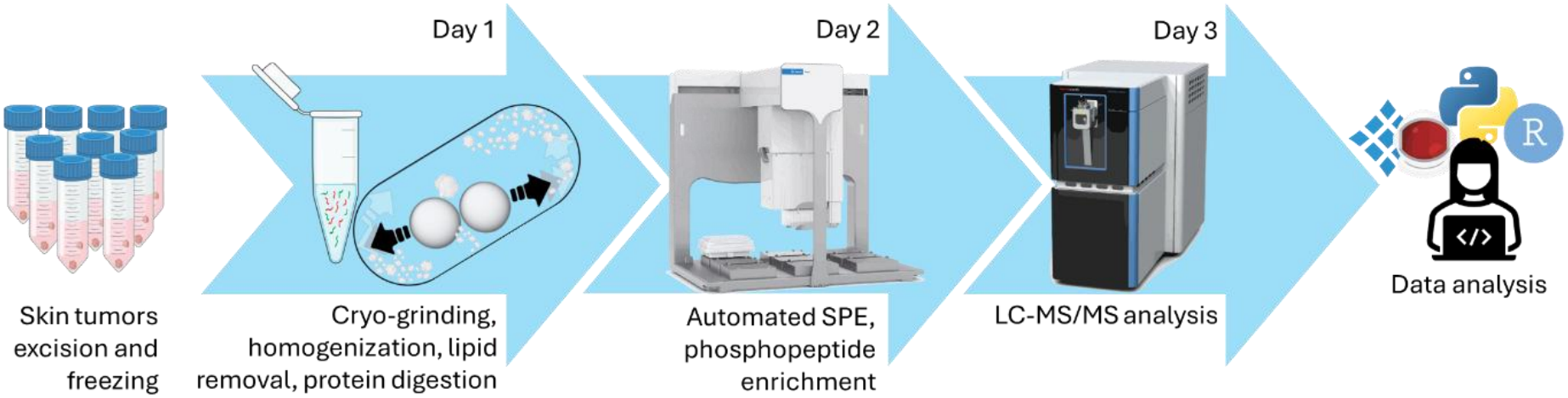

## Before you begin

Skin is the outermost organ of the human body and the first line of defense against trauma, infections and radiation^1^.More than 3,000 diseases affect the skin, varying in severity and prevalence and impacting individuals of all ages and social backgrounds^1–3^ Skin-related conditions are a global burden for public health, as nowadays they represent the most common reason to consult a general practitioner^4^. Some of these diseases, such as skin cancer, can be life threatening, many more significantly affect the quality of life of affected individuals^3^. Understanding skin homeostasis and pathology is essential to improve diagnosis and treatment. Mass spectrometry (MS)-based proteomics represent a powerful tool for obtaining a global and unbiased view of the molecular players^2^. However, the high abundance of lipids in skin specimens and the presence of crosslinked proteins make skin a challenging tissue to analyze^1,3,5^. Here, we present aworkflow for deep proteomic and phosphoproteomic profiling of snap-frozen biopsies, enabling the quantification of up to 8,000 proteins and up to 60,000 phosphosites.

### Innovation

Other sample preparation workflows for human skin focus on downstream proteome measurement and rely on detergent-based^1,6^ or acid-aided lysis,^5^ or direct digestion^7^. Phosphoproteomic analyses are rarely performed and workflows are not tested for compatibility. In contrast, our approach enables both deep proteomic and phosphoproteomic analyses from snap-frozen skin biopsies using a single streamlined sample preparation pipeline. The integration of cryogenic tissue disruption and chloroform–methanol precipitation ensures efficient protein extraction and contaminant removal while maintaining compatibility with downstream phosphopeptide enrichment, expanding skin proteomics beyond protein-level analyses.

### Institutional permissions

Ethical approval should be obtained before the acquisition of patient or animal data. It should be requested early in the planning process.

For the development and validation of this workflow, surplus skin samples were obtained from routine surgeries at the Dermatology Department of the University Hospital Zürich with the assistance of the SKINTEGRITY.CH biobank (PB_2018-00194)). Informed consent had been obtained from all patients and all experiments conformed to the principles set out in the WMA Declaration of Helsinki and the Department of Health and Human Services Belmont Report. The use of material for research purposes was approved by the cantonal ethic commission of Bern (KEK Bern; Project-ID 2022-02278).

## Key resources table

| REAGENT or RESOURCE | SOURCE | IDENTIFIER |
|---|---|---|
| Chemicals, peptides, and recombinant proteins |  |  |
| Sodium chloride | Roth | 3957.4 |
| Sucrose | Sigma-Aldrich | 84097 |
| EDTA (ethylene diamine tetra acetate) disodium salt | Sigma-Aldrich | E5134 |
| Trifluoroacetic acid (TFA) | Sigma-Aldrich | 302031 |
| MS-grade Water | VWR | 23595.328 |
| MS-grade Acetonitrile | VWR | 20060.32 |
| Ethanol | N/A | N/A |
| Benzonase | Dr. Nuclease Benzonase | BP4200 |
| Sequencing grade modified trypsin | Promega | V5113 |
| Dithiothreitol | VWR | 0281 |
| Iodoacetamide | Sigma-Aldrich | 4386 |
| Water with 0.1 % Formic Acid (v/v), Optima LC/MS grade | Fisher Scientific | LS118 |
| Acetonitrile with 0.1 % Formic Acid (v/v), Optima LC/MS grade | Fisher Scientific | LS120 |
| Ammonium bicarbonate | Sigma-Aldrich | 9830 |
| Urea | Sigma-Aldrich | U5378 |
| Trizma | Sigma-Aldrich | T1503 |
| PhosSTOP tablets | Roche | 4906837001 |
| Complete ™ EDTA-free protease inhibitor | Roche | 5056489001 |
| Acetone | Sigma-Aldrich | 270725 |
| Lysyl Endopeptidase | IGZ Instruments | 129-02541 |
| Methanol | Merck | 1060354000 |
| Chloroform | VWR | 83627.2909 |
| Software and algorithms |  |  |
| Spectronaut | Biognosys | N/A |
| Protein Sample Prep Workbench Software | Agilent | Phosphopeptide Enrichment v2.1 |
| Other |  |  |
| Bravo liquid handling platform (Bravo) | Agilent | RRID:SCR_026137 |
| Orbitrap Astral mass spectrometer | Thermo Fisher Scientific | RRID:SCR_026205 |
| Vanquish Neo UHPLC system | Thermo Fisher Scientific | RRID:SCR_026495 |
| Greiner Bio-One 96-well microplate for PCR, skirted | Greiner | 652270B |
| NanoDrop One Microvolume UV-Vis Spectrophotometer | Thermo Scientific | ND-ONE-W |
| Omni 2mL Stainless Steel Cryo Tubes | Omni | OMNI-19-6001 |
| Mixer mill MM400 | Retsch | RRID:SCR_020427 |
| AssayMAP 25 µL Reversed Phase (RP-S) Cartridges | Agilent | G5496-60023 |
| Custom machined aluminium 2 ml tubes adapters for retsch | N/A | N/A |
| 5 mm stainless steel beads | Fisher scientific | 17211762 |
| 1 ml deep 96-well plate | Agilent | 201276-100 |
| 50 ml testing tubes | N/A | N/A |
| Screw caps for micro tubes | Sarsted |  |
| Reservoir, 12 column, polypropylene, 7 mL/column | Agilent | 201280-100 |
| Multichannel pipet | N/A | N/A |
| AssayMAP Cart Rack, Fe(III)-NTA cartridges | Agilent | G5496-60085 |
| Eppendorf ThermoMixer® C | Eppendorf | 5382000015 |
| ThermoTop® | Eppendorf | 5308000003 |
| Orbital shaking station | Agilent | G5431B |
| Bravo 96 AssayMAP Head | Agilent | G5058A |
| PCR strips and lids | N/A | N/A |
| 1.5 ml testing tubes | N/A | N/A |

## Materials and equipment

### Grinding equipment setup

The grinding tubes were modified by replacing the standard rubber gasket with a PTFE (Teflon) gasket to improve physical resistance and maintain sealing integrity under cryogenic conditions (Figure 1A).

**Figure 1:**
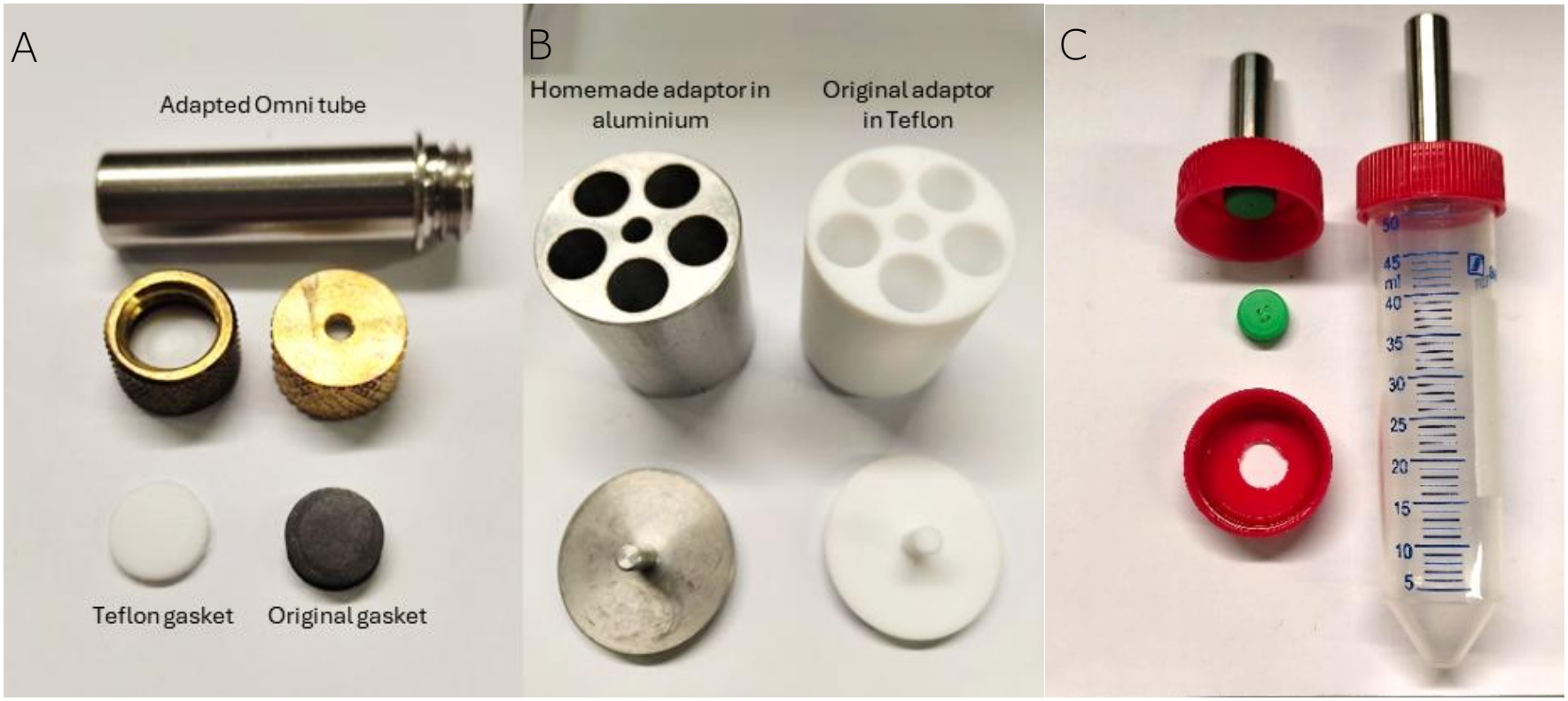
Grinding and sample recovery setup. A. Adapted omni metal tubes with PTFE-based gasket. B. custom-machined aluminium adapter. C. Lysate recovery setup.

In the original PTFE-based 2 ml tube adapter system, cryogenic cooling led to differential thermal contraction, resulting in the metal tubes becoming tightly fitted and difficult to remove after grinding. To overcome this limitation, the adapter was replaced with a custom-machined aluminium version, which has a reduced contraction at liquid nitrogen conditions (Figure 1B).

### Sample recovery components

To facilitate recovery of lysate while minimizing bead carryover, a centrifugation-assisted collection setup was implemented. The metal grinding tube was positioned inverted and fixed to a modified 50 mL Falcon tube cap containing an opening, allowing the lysate to be transferred through the tube under centrifugal force while retaining beads in the grinding chamber. This approach reduces the need for pipetting-based recovery steps, which can be inefficient with viscous or particulate-containing lysates (Figure 1C).

#### Homogenization buffer

| Reagent | Final concentration |
|---|---|
| Tris (Trizma) pH 7.5 | 250 mM |
| Sucrose | 290 mM |
| EDTA | 25 mM |

Can be autoclaved for 6 months storage at 20-25°C. Add cOmplete and PhosSTOP inhibitors at manufacturer concentration before use.

#### Urea buffer

| Reagent | Final concentration |
|---|---|
| Tris (Trizma) pH 7.5 | 100 mM |
| Urea | 8 M |

Prepare fresh before usage. Add cOmplete and PhosSTOP inhibitors at manufacturer concentration.

#### Dithiothreitol (DTT) stock solution

| Reagent | Final concentration |
|---|---|
| DTT | 100 mM |
| ddH<sub>2</sub>O | - |

Storage at -20°C 2-3 months.

#### Iodoacetamide (IAA) stock solution

| Reagent | Final concentration |
|---|---|
| IAA | 550 mM |
| ddH<sub>2</sub>O | - |

Storage at -20°C in dark tubes 2-3 months.

#### ABC buffer

| Reagent | Final concentration |
|---|---|
| Ammonium bicarbonate | 100 mM |
| ddH<sub>2</sub>O | - |

Storage at 20-25°C 2-3 months.

#### Acetone 80%

| Reagent | Final concentration |
|---|---|
| Acetone | 80% v/v |
| ddH<sub>2</sub>O | - |

Storage at -20°C 6 months.

#### TFA 5%

| Reagent | Final concentration |
|---|---|
| Trifluoroacetic acid | 5% v/v |
| ddH<sub>2</sub>O | - |

Storage at 20-25°C 6 months.

### Automated Solid-Phase Extraction (SPE)

#### Prime and Elution buffer

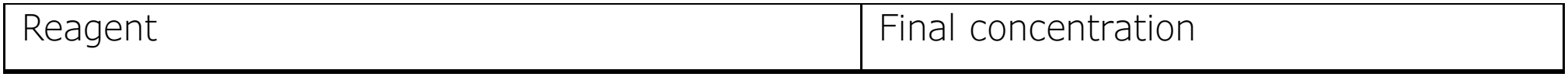
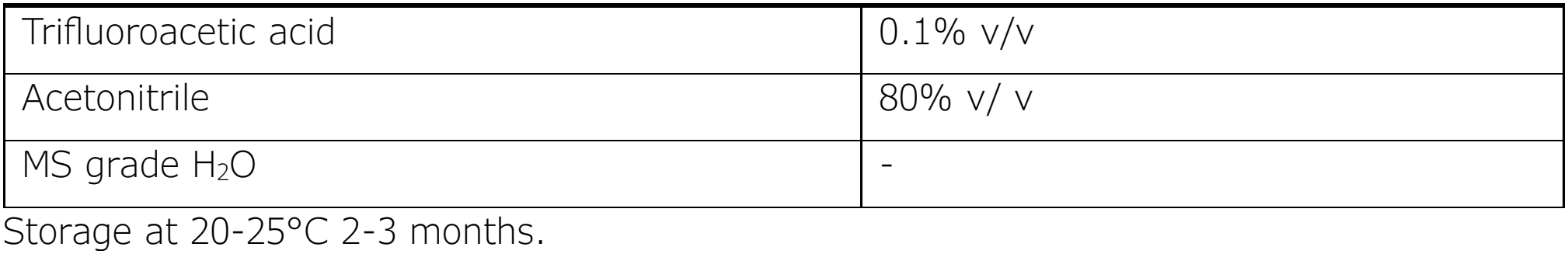

#### Equilibration, Wash and Cartridge Wash buffer

| Reagent | Final concentration |
|---|---|
| Trifluoroacetic acid | 0.1% v/v |
| MS grade H<sub>2</sub>O | - |

Storage at 20-25°C 2-3 months.

### Phospho-enrichment

#### Equilibration, Wash and Cartridge Wash Buffer

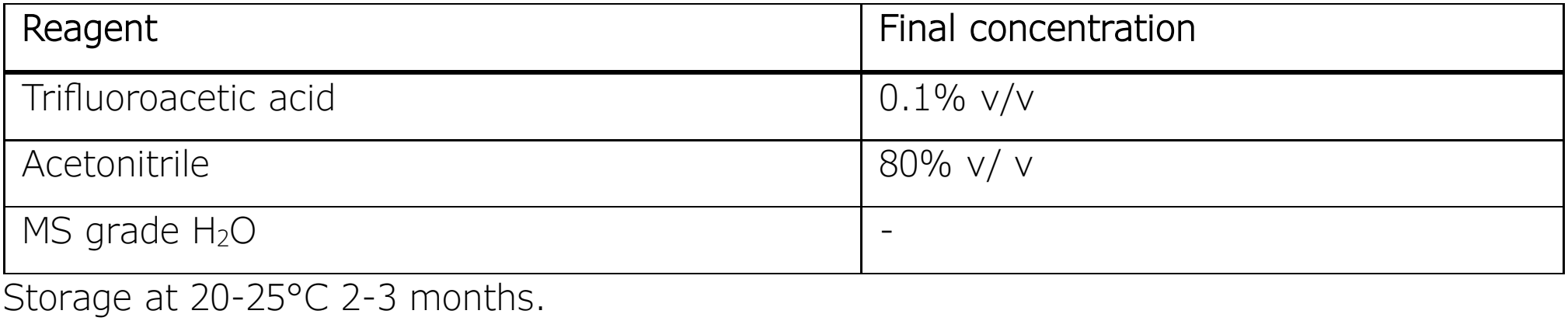

#### Prime Buffer

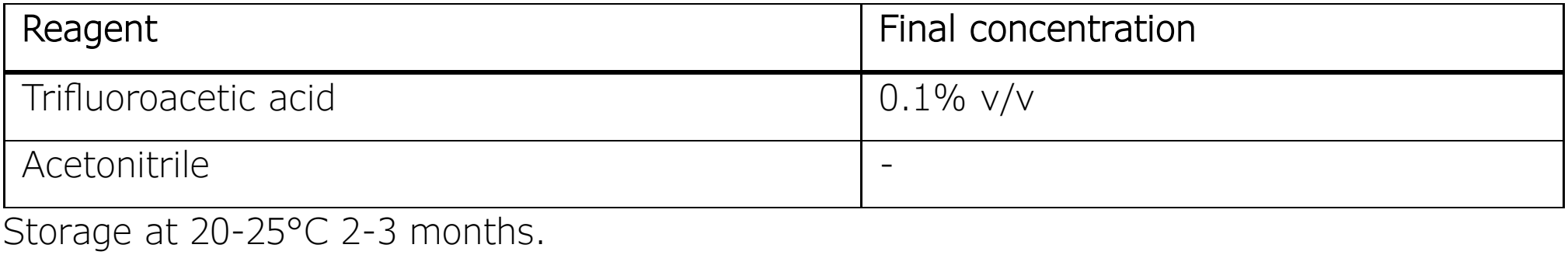

#### Elution Buffer 1

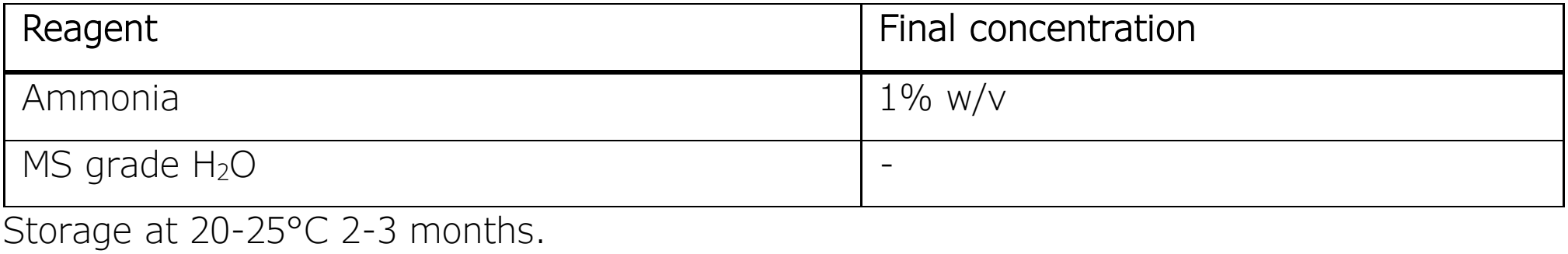

#### Elution Buffer 2

| Reagent | Final concentration |
|---|---|
| Ammonia | 1% w/v |
| Acetonitrile | 80% v/v |
| MS grade H<sub>2</sub>O | - |

Storage at 20-25°C 2-3 months.

## Step-by-step method details

### Cryogenic Pulverization

The following steps ensure effective cryogenic grinding of tissue samples preventing thawing using an in-house setup.

Timing: ~1 h

**Note:** While this protocol describes implementation using the Retsch MM400 system with a custom adapter, cryogenic pulverization can also be achieved using alternative commercially available milling or homogenization platforms suitable for cryogenic operation.

1. Pre-cool adapted Omni metal tubes, homemade metal adapters, and beads until they are at liquid nitrogen temperature

#### CRITICAL

Keep the tubes cold (LN or −80°C) until homogenization buffer is added to prevent thawing of the samples (Troubleshooting 1)

2. Transfer tissue samples (~2 mm^3^) into pre-cooled tubes.
3. Add 2 pre-cooled beads to each tube.
4. Place tubes into the pre-cooled metal adapter.
5. Perform mechanical disruption cycle at 27 Hz for 1 min with Retsch MM400.
6. After each cycle re-cool the tubes and adapters in liquid nitrogen to keep the sample frozen.
7. Repeat for a total of 10 cycles (Troubleshooting 2):

**Pause point:** the tubes containing cryo-grinded tissue can be stored at −80°C.

### Homogenization of samples

This step homogenizes cryogenically disrupted tissue in lysis buffer to enable efficient protein extraction for downstream proteomic and phosphoproteomic analyses.

Timing: 20 min

8. From this step forwards, cooling is no longer needed
9. Spin the tubes in a table centrifuge for 30 sec to collect the grinded sample at the bottom.
10. Add 500 µl of homogenization buffer (250 mM Tris, 250 mM sucrose, 25 mM EDTA with freshly supplemented PhosSTOP and cOmplete tablet) to each tube. **Note:** If the homogenization buffer freezes in the tubes, let it thaw at room temperature before proceeding.
11. Flick the tube once and spin to recover powdered sample eventually stuck in the cap
12. Exchange the cap to a Sarstedt plastic one to avoid leaks
13. Shake at 20 Hz for 6 minutes to homogenize the samples.

### Centrifugation-assisted homogenate recovery

This setup enables efficient separation of beads from the homogenate and facilitates lysate recovery while minimizing sample loss.

Timing: 5 min

14. Use the custom 50 ml-centrifugation tube setup to recover the homogenate from the tubes **Note:** the sample can be recovered by pipetting, however the loss of samples will be higher.
15. Centrifuge samples at 125 rcf for 2 min to collect the homogenate

### Chloroform/Methanol precipitation

Chloroform allows the elimination of hydrophobic contaminants such as lipids from the samples while methanol precipitates the proteins.

#### CRITICAL

this step should be carried out in a fume hood Timing: 1 h

16. Add 500 µl of chloroform and 1500 µl of water to the homogenate
17. Vortex
18. Centrifuge at 4500 rcf for 10 min
19. Remove carefully the aqueous layer (top one)
20. Add 1500 µl of Methanol
21. Vortex
22. Centrifuge at 4500 rcf for 10 min
23. Remove the supernatant (Troubleshooting 3)
24. Add 1 ml of 80% acetone and transfer the samples to 2 ml tubes **Note:** the sample can be transferred with a p1000 pipette.
25. Centrifuge 21,000 rcf for 10 min
26. Remove the supernatant
27. Wash twice with 80% acetone
28. Remove supernatant
29. Let the samples dry for 10 min

**Pause point:** the precipitated proteins can be stored at −80°C.

### Resuspension and digestion

In this step precipitated proteins are resuspended in denaturing buffer and enzymatically digested into peptides suitable for bottom-up LC-MS/MS analysis.

Timing: 3 h + 16 h

30. Resuspend the samples in 8 M urea supplemented with PhosSTOP (Troubleshooting 4)
  a. Use 2 times the pellet volume (50-100 µl) of urea buffer

#### CRITICAL

The maximum volume must not be more than 100 µl, otherwise the sample will not fit into the plate for the automated sample cleanup.

**Note:** Sonicate if necessary.

31. Add DTT to 1 mM final concentration
32. Incubate 15 min at 25°C, shaking at 650 rpm
33. Add IAA to 5.5 mM final concentration
34. Incubate 15 min in the dark at 25°C, shaking at 650 rpm

### Add 3 µg of LysC (Potential solution

The protein pellet was too dry, or the resuspension buffer volume was too low. Resuspend the pellet in an appropriate volume of resuspension buffer and assist solubilization by brief sonication in a water bath or by pipetting.

35. Troubleshooting 5)
36. Incubate 2 h at 25°C shaking at 650 rpm
37. Dilute the sample to ≤ 1 M Urea with 100 mM Tris pH 8 (Troubleshooting 2)

### Add 1:100 trypsin (Potential solution

The protein pellet was too dry, or the resuspension buffer volume was too low. Resuspend the pellet in an appropriate volume of resuspension buffer and assist solubilization by brief sonication in a water bath or by pipetting.

38. Troubleshooting 5)
39. Incubate 16 h at 25°C

### Acidification

This step is needed to acidify the sample before clean up, charging all peptides.

40. Acidify to 2% TFA
41. Centrifuge at maximum speed for 10 min to pellet eventual residues
42. Load supernatant into 96-deep-well plate for Turbo SPE **Note:** The robot works column by column

### Automated SPE

This step allows peptide clean-up and desalting to support phosphopeptide enrichment. Timing: 3-5 h

43. Place the Reverse-Phase cartridges in the matching positions occupied by the samples in the 96-deep-well plate. **Note:** Add resin-free cartridges to unoccupied spots in respective columns for correct pressure handling.
44. Use the Peptide Cleanup v3.0 application for the Bravo liquid handling platform
45. Place the metal rack with the cartridges in position 2 (Figure 2A)
46. Add the flow-through 96-deep well plate to position 7 (Figure 2A)
47. Add eluate collection U-bottom 96-well plate to position 9 (Figure 2A)
48. Fill up the necessary columns of the priming, equilibration and elution reservoir plates with the corresponding buffers (in positions 5, 8 and 6 of the robot, respectively – Figure 2A).
49. Place the plate containing the samples in position 4 (Figure 2A)
50. Add the plate to collect the organic waste in position 3 (Figure 2A).
51. Set the software as follows:
  a. Set “Number of Full Columns of Cartridges” to the number of columns in the experiment.
  b. Adjust the labware according to what is used
  c. Tick the boxes for:
    i. “Prime” → 250 µl – 300 µl/min – 1 wash cycle.
    ii. “Equilibrate” → 250 µl – 10 µl/min – 0 wash cycle.
    iii. “Load Samples” → 800 µl – 5 µl/min – 0 wash cycle.

**Figure 2:**
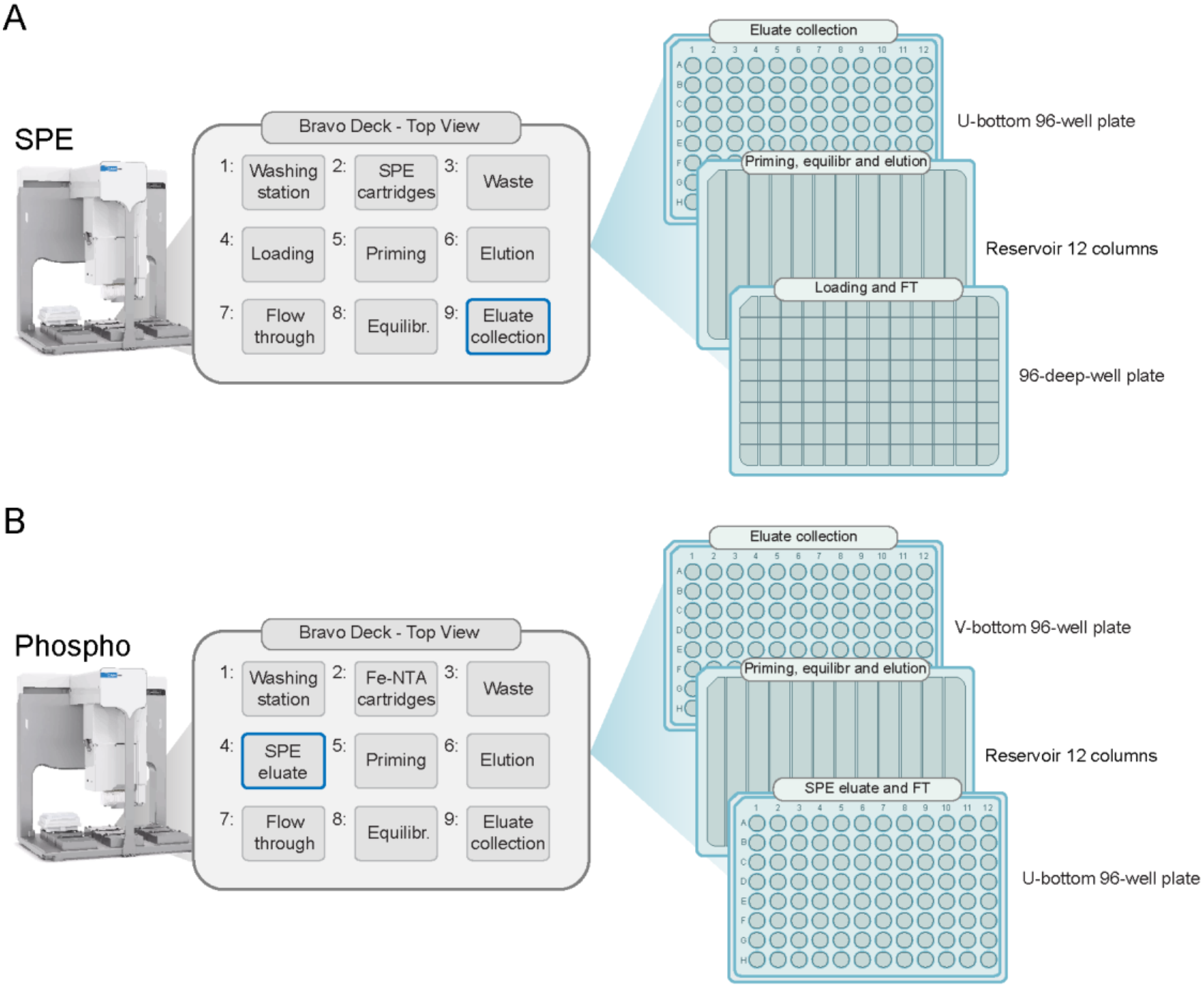
Setup for the bravo liquid handling platform deck for A. Solid-Phase Extraction (SPE) and B. phosphopeptides enrichment. Rows are labeled A-H, columns 1-12.

#### CRITICAL

The volume of this step depends on the volume of the samples. 800 µl is the recommended maximum.

iv. “Collect Flowthrough”.
v. “Internal Cartridge Wash” → 250 µl – 10 µl/min – 0 wash cycle.
vi. “Elute” → 230 µl – 5 µl/min – 2 wash cycle.

52. Run the protocol **Note:** The time depends on the volume of the samples. With 800 µl of samples the protocol takes around 4 h 30 min.
53. At the end of the clean-up protocol, store 20 µl of the eluate in the PCR strip. It can be stored at −80°C or loaded directly for MS-based proteomics.

### Phosphopeptide-enrichment

This step selectively enriches phosphopeptides from the peptide mixture, enhancing detection of low-abundance phosphorylation events during LC-MS/MS analysis.

Timing: 2 h 30 min

54. Place the Fe-NTA cartridges to the matching position occupied by the eluates in the U-bottom 96-well plate.
  a. Keep the resin-free cartridges from before
55. Use the Phospho-peptide Enrichment v2.1 application for the Bravo liquid handling platform
56. Place the metal rack with the cartridges in position 2 (Figure 2B)
57. Add the flow through collection plate in position 7 (Figure 2B)
58. Fill up completely the necessary columns of the priming and wash reservoir plates with the corresponding buffers (in positions 5 and 8 of the robot, respectively – Figure 2B).
59. Place the plate containing the SPE eluates in position 4 (Figure 2B)
60. Add a plate to collect the organic waste in position 3 (Figure 2B)

#### CRITICAL

Do not fill the elution buffer reservoir plate yet.

61. Set “Number of Full Columns of Cartridges” to the number of columns in the experiment.
62. The phosphopeptide-enrichment is done in multiple rounds:
  a. Loading
    i. Tick the boxes for
      1. “Prime” → 100 µl – 300 µl/min – 1 wash cycle.
      2. “Equilibrate” → 200 µl – 10 µl/min – 1 wash cycle.
      3. “Load Samples” → 200 µl – 5 µl/min – 3 wash cycles.
      4. “Collect Flowthrough”.
    ii. Click “Run Protocol” **Note:** This phase takes around 60 min **Note:** At the end of this phase, collect non-phosphorylated flowthroughs in a PCR strip and freeze them at −80°.
  b. Washing
    i. Untick all the previous boxes
    ii. Fill up the Equilibration/Internal Cartridge Wash Buffer reservoir
    iii. Tick the box “Internal Cartridge Wash”
    iv. Click “Run Protocol” **Note:** This phase takes 20 min
  c. Washing and First Elution
    i. Fill the necessary columns in the reservoir plate for Elution Buffer 1
    ii. Place it in position 6 (Figure 2B)
    iii. Add 5 µl of 100% formic acid to the wells of the eluate collection v-bottom 96-well plate
    iv. Place it in position 9 (Figure 2B)
    v. Keep the “Internal Cartridge Wash” ticked
    vi. Tick the box “Elute” → 50 µl – 5 µl/min – 1 wash cycle
    vii. Click “Run Protocol” **Note:** This phase takes 30 min
  d. Second elution
    i. Fill the necessary columns in the reservoir plate for Elution Buffer 2
    ii. Exchange it with the other Elution plate in position 6 (Figure 2B)
    iii. Untick everything except for “Elute”
    iv. Click “Run Protocol” Note: This phase takes 10 min
63. Collect the elution plate
  i. Move the samples to PCR strips
  ii. Freeze them at −80°C
64. Dry the phospho-& non phosphorylated samples in freeze dryer overnight
65. Resuspend in 20 µl of 0.1% formic acid for LC-MS/MS analysis.

## Expected outcomes

This workflow is expected to generate peptide samples compatible with whole proteome and phosphoproteome LC-MS/MS analyses from cryopreserved tissue specimens. Efficient cryogenic disruption combined with protein precipitation should yield reproducible protein recovery across samples while minimizing common sources of variability such as contaminants and missed cleavages. The number of quantified proteins and phosphosites depends on the quality of the sample and on the mass spectrometer used. When using the newest generation of instruments and snap frozen material, we expect that data independent acquisition can quantify up to 8,000 proteins and up to 60,000 phosphosites. GO term analysis on the identified proteins highlights the broad coverage of the proteome (Figure 3Figure 3).

**Figure 3:**
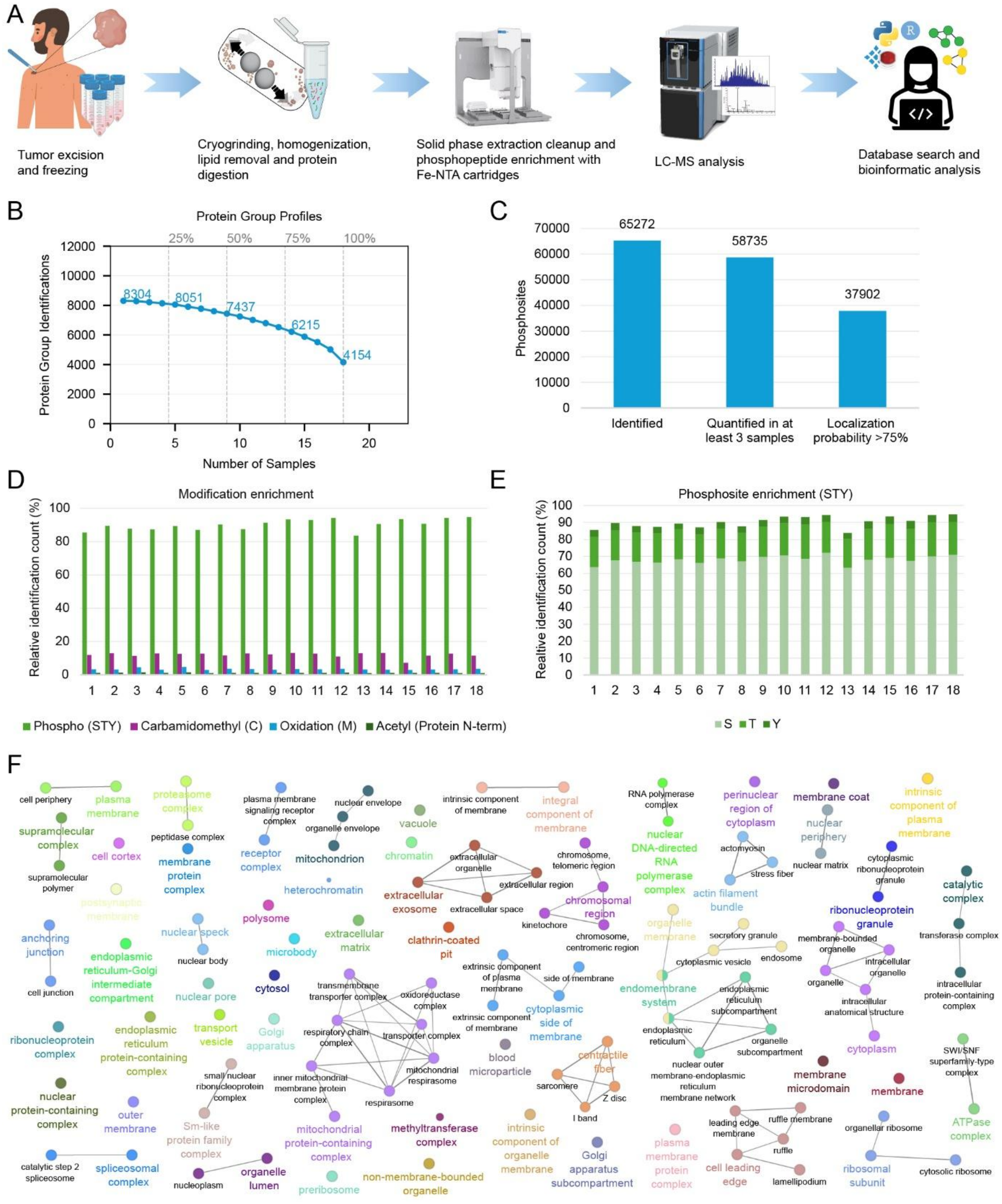
Overview of the proteomics workflow and quality control metrics for skin (phospho)proteomics. A. Schematic representation of the sample preparation workflow showing the streamlined process to obtain both proteome and phosphoproteome from the same sample. B. Protein group profiles showing the identification of proteins over 18 biopsies of healthy and diseased skin. C. Bar charts depicting the number of phosphosites quantified across 18 biopsies of healthy and diseased skin, reflecting the efficiency of phosphopeptide enrichment and the depth of phosphoproteome coverage. Class I phosphosites are defined based on maximum localization probability of ≥75%. D. Relative percentage distribution of different post-translational modifications (PTMs) identified in the dataset, highlighting the enrichment specificity. E. Relative percentage distribution of phosphorylated residues (serine, threonine, and tyrosine), demonstrating the expected predominance of serine and threonine phosphorylation and the extent of tyrosine phosphorylation. F. Gene Ontology (GO) Cellular Component enrichment analysis of the identified proteome, showing broad subcellular coverage including membrane complexes, cytoplasm, mitochondria, extracellular space, and nuclear compartments. Results were created and visualized with ClueGO v2.5.10^8^.

## Troubleshooting

### Troubleshooting 1

Sample does not grind or remains stuck to beads. The tissue may have partially thawed during transfer, or beads may have warmed causing the sample to adhere to the tube walls rather than fracturing upon impact.

#### Potential solution

Ensure all tubes, beads, and tube racks are pre-chilled in liquid nitrogen immediately before loading the sample in the pre-chilled adapter. Work as rapidly as possible during tissue transfer to avoid any warming. Verify that the bead is moving freely before starting the run by briefly hitting the frozen tube against a hard surface.

### Troubleshooting 2

Insufficient tissue disruption after grinding (visible tissue fragments remain).

### Potential solution

The grinding cycle was too short, the oscillation frequency was too low, or the bead-to-tissue ratio was suboptimal. Increase the number of grinding cycles or extend the duration of each cycle. Re-submerge tubes in liquid nitrogen between cycles to prevent thawing. Ensure the correct bead size is used for the tissue type.

Troubleshooting 3

Low protein yield after precipitation.

### Potential solution

Incomplete precipitation due to insufficient incubation time or temperature, or loss of pellet during supernatant removal. Ensure the correct ratio between chloroform and methanol is used. When removing the supernatant, leave a small residual volume to avoid disturbing the pellet.

### Troubleshooting 4

Poor or inconsistent protein resuspension after precipitation.

### Potential solution

The protein pellet was too dry, or the resuspension buffer volume was too low. Resuspend the pellet in an appropriate volume of resuspension buffer and assist solubilization by brief sonication in a water bath or by pipetting.

### Troubleshooting 5

High proportion of missed cleavages detected in the final dataset.

### Potential solution

Insufficient dilution of the sample prior to trypsin digestion leads to elevated concentrations of urea that inhibit trypsin activity, resulting in incomplete digestion and a high missed cleavage rate. Ensure the sample is diluted to <1 M urea before adding trypsin. Increase LysC- or trypsin-to-protein ratios.

## Limitations

This protocol was optimized for tissue samples of approximately 2 mm^3^. Larger samples may not be efficiently processed within the grinding tubes and can impair efficiency. Therefore, tissue should be pre-cut into smaller fragments prior to cryogenic grinding to ensure consistent disruption and prevent material retention in the tube.

This protocol was implemented using a cryogenic grinding setup based on the Retsch MM400 with a custom adapter system. While this configuration provides robust tissue disruption under cryogenic conditions, alternative cryogenic milling or homogenization platforms may be required depending on instrument availability and sample type.

## Resource availability

Lead contact: Prof. Joern Dengjel

Technical contact: Sibilla Sander

## Acknowledgments

This work was supported by the University and the Canton of Fribourg as part of the SKINTEGRITY.CH research network, the Swiss National Science Foundation with the NRP79 (grant 407940_206394).

## Author contributions

Conceptualization: S.S., M.S., I.B., J.D.; Methodology: S.S.; Validation: S.S., I.B.; Formal analysis: S.S., I.B.; Investigation: S.S., I.B.; Resources: G.R, M.L.; Writing - Original Draft: S.S., J.D., Writing - Review & Editing: S.S., I.B., M.S., G.R, M.L, J.D., Visualization, S.S., Supervision: J.D.; Project administration: J.D.; Funding acquisition: J.D.

